# Avian telencephalon and cerebellum volumes can be accurately estimated from digital brain endocasts

**DOI:** 10.1101/2024.12.04.626912

**Authors:** Aubrey R. Keirnan, Felipe Cunha, Sara Citron, Gavin Prideaux, Andrew N. Iwaniuk, Vera Weisbecker

## Abstract

For studies of the evolution of vertebrate brain anatomy and potentially associated behaviours, reconstructions of digital brain endocasts from computed tomography scans have revolutionised our capacity to collect neuroanatomical data. However, measurements from digital endocasts must be validated as reflecting actual brain anatomy, which is difficult because the collection of soft tissue information through histology is laborious and time consuming. In birds, the reliability of digital endocast measurements as volume proxies for the two largest brain regions – the telencephalon and cerebellum - remains to be validated despite their use as proxies e.g. of cognitive performance or flight ability. We here use the largest dataset of histology and digital endocasts to date, including 136 species from 25 avian orders, to compare digital endocast surface area measurements with actual brain volumes of the telencephalon, cerebellum, and whole-brain endocast. Using linear and phylogenetically informed regression analyses, we demonstrate that endocast surfaces are strongly correlated with their brain volume counterparts for both absolute and relative size. This provides empirical support for using endocast-derived cerebellar and telencephalic surface areas in existing and future studies of living and extinct birds, with potential to expand to the dinosaur-bird transition in the future.

## INTRODUCTION

Over the past two decades, digital endocasts have become an increasingly important tool in the study of neural and behavioural evolution in vertebrates [1–6]. Using computed tomography (CT) scanned skulls, the brain cavity is digitally “filled” to approximate the volume and shape of the brain [7]. These methods are largely non-destructive, allowing research into the neuroanatomy of rare [8] and extinct species [9], including fossil specimens [10, 11]. For preservation purposes, such as museum specimens or fossils embedded in sedimentary matrix, CT scanning offers non-destructive access to otherwise unattainable information [12, 13]. To date, endocasts have been used to study brain evolution across all vertebrate classes, including fishes [5], amphibians [2], birds [14], non-avian reptiles [15, 16], and mammals [6, 17, 18]. Therefore, our understanding of the brain-to-endocast relationship is fundamental to the study of vertebrate brain evolution.

Our knowledge of evolutionary patterns in brain size [13, 19], shape [6, 20], and composition [21] has advanced substantially over the past century through endocast comparisons [13, 22]. For example, early endocast research on pterosaur fossils, an extinct clade of flying reptiles, revealed similarities between their brains with those of modern birds [23]. These included the positions of the optic lobes (or midbrain) and the cerebellum, distinguishing pterosaurs and birds from other reptiles [23]. More recently, endocast studies have been used to infer brain–behaviour relationships, including social behaviour in big cats [24], nocturnality in birds [8, 9, 25], and the habitats of snakes [16]. However, there is substantial variation in the brain–endocast relationships among vertebrates [26], yet the use of endocast measurements relies on the assumption that they are indicative of the brain region of interest [2, 3, 12]. While some studies have verified the brain–endocast correlations for particular brain regions [27, 28], measurement modalities such as shape or volume are not necessarily interchangeable [4, 6], meaning that both must be tested to establish the validity of these methods. For fishes, amphibians, and non-avian reptiles, in which brains do not fill the cranial cavity, inferences are limited as endocasts overestimate brain size [2, 4, 29]. This can be an issue particularly at the root of iconic radiations such as dinosaurs, whose closest living non-avian relatives – the crocodiles – have brains that typically occupy less than half of their endocranial space in adults [26, 30]. In contrast, the endocranial volume in birds is strongly correlated with brain volume [31].

The brains of birds have been the subject of extensive efforts to establish the relationship between the endocranial cavity and brain tissue [22, 28, 31–33]. Their endocasts are remarkably faithful to the shape of the brain itself, and endocranial volume is a valid proxy for actual brain volume and widely used in studies of brain size (e.g. [14, 31, 34, 35]). Further, several brain regions are discernible in endocasts [10, 12, 22, 28]. The endocast surface areas of two of these brain regions, the wulst and optic lobes, are reliable estimates of the underlying hyperpallium and optic tectum regions, respectively, of the brain [28]. However, inferences have also been drawn from endocast measurements of the telencephalon and cerebellum, even though the reliability of measuring these brain regions from endocasts remains untested. Changes in the size of both regions are important in the evolution of differences in avian cognition [36] and the brain becoming “flight-ready” [37, 38]. For example, the expansion of the telencephalon is a key anatomical change in the evolution of the avian brain [37, 39] and associated with higher rates of innovative behaviour [40]. Similarly, changes in relative cerebellum size, and part of the cerebellum, are associated with behavioural differences across avian clades [41, 42]. They also comprise the majority of brain volume [43] and the endocast surface. Exploiting the potential of these two brain regions in explaining major transitions in avian brain evolution therefore requires validation of how faithfully their endocast imprints reflect their soft tissue volume.

Here, we determine the brain–endocast relationships of the telencephalon and cerebellum using the largest, phylogenetically diverse sample of bird species to date. While brain regions are typically measured as volumes, endocasts lack the internal anatomical landmarks needed to accurately divide them into volumes. This is especially true of the avian cerebellum, because its anterior surface is covered by the posterior telencephalon. While surface area may not be interchangeable with volume [6], we hypothesised this comparison would show a typical isometric relationship, close to or equal to a 2:3 ratio or surface area increasing at a rate of 0.67 to that of the volume. However, as the relative size of these brain regions is often important for studying neurobehavioural evolution [3, 26, 32, 44, 45], we also asked if their relative size when measured from endocasts was comparable to that of brains. We expected that, if the endocast reliably estimates the relative size of these regions, they would produce comparable relationships, which would result in similar slopes.

## METHODS

### Specimens

Data on whole brain, telencephalon and cerebellum volumes were compiled from the literature [41, 43, 46–56] and supplemented with previously unpublished data from a histological collection maintained by ANI [57]. These specimens were provided to ANI dead and were not euthanised for the purpose of this study. Further, all procedures were completed in accordance with the guidelines and policies of the Canada Council on Animal Care. All of these brains were gelatin-embedded and serially sectioned in the coronal plane at a thickness of 40µm. After mounting and staining with thionin acetate, the volumes were measured using unbiased stereology in StereoInvestigator (Microbrightfield Inc., VT, USA) by FC. For some species, the telencephalon and cerebellum volumes were measured from different individuals. In such cases, we included the brain volumes for both individuals to accurately calculate the relative size of brain regions. More details on the brain processing and volume measurement methods are described in [46]. All data and their specific sources are provided in the electronic supplementary material [58].

To compare the brain volumes with surface areas from endocasts, we collected existing micro-computed tomography (μCT) scans as well as completing additional scans for this study. Existing scans were sourced through the online repository Morphosource (Project ID: 000642669), our previous studies [8, 25] as well as completing additional scans. The source, museum identification, and the scanning parameters for each specimen are provided in Supplementary Table 1. In total, we had brain volume measurements and CT scans from 136 species, representing 58 families and 25 orders. However, for four species, the volumes for either the telencephalon or cerebellum was unavailable – two species from each region - reducing the total to 134 species for these brain regions.

### Endocast reconstruction and surface area data collection

All endocast reconstruction and measurements were completed by AK. We imported the CT scans into the image processing software Materialise Mimics (v.24.0; released in 2021 by Materialise NV) and exported reconstructed skulls. Meshes of the inner surface of the brain cavity – the endocast – were derived using the endomaker function from the Arothron package [59] in R [60] with the meshes decimated to 20000 but otherwise using the base settings for all reconstructions. The meshes were incomplete due to natural openings or damage to the skull, and often included excess material. To complete the endocasts, these were imported into Geomagic Wrap (v.2021.2.2; released in 2023 by 3D Systems) to remove excess material and patch openings using the “flat” fill setting. Bumps and crevices were also removed and subsequently patched with the “flat” fill along with the removal and patching of nerves and blood vessels, following best practices as outlined by Balanoff, Bever [33] (example in electronic supplementary material figure S1). As is to be expected, there was some variation in the amount removed and patching needed for each specimen. However, efforts were made to keep the reconstructions consistent and maintain the surface details of all the endocasts for accuracy.

We defined the borders for the telencephalon and cerebellum by consulting previous studies which partitioned endocasts for volume, surface area or landmark based measurements [4, 8, 21, 28, 61]. The posterior border of the telencephalon fossa was defined as the junction where it meets the cerebellar fossa and continues ventrally through the crista tentorialis above the fossa tecti mesenchephali. These borders then meet rostrally above the optic chiasma. For the cerebellum fossa, the dorsal border was defined by the junction that meets the telencephalon fossa and continues along the caudal border of the fossa tecti mesencephali on either side. This border continues as the crista present at the ventral region of the cerebellum fossa, dorsal to the brainstem fossa, meeting in the posterior region of the foramen magnum. We used the surface areas within these borders, which encompass the visible imprints of both regions to estimate their sizes from the endocasts (Figures 1a, b). We collected all measurements using Geomagic Wrap by following the borders outlined above. In Figure 1c, we provide examples of these selections from each of the 25 clades in this study.

**Figure 1.**
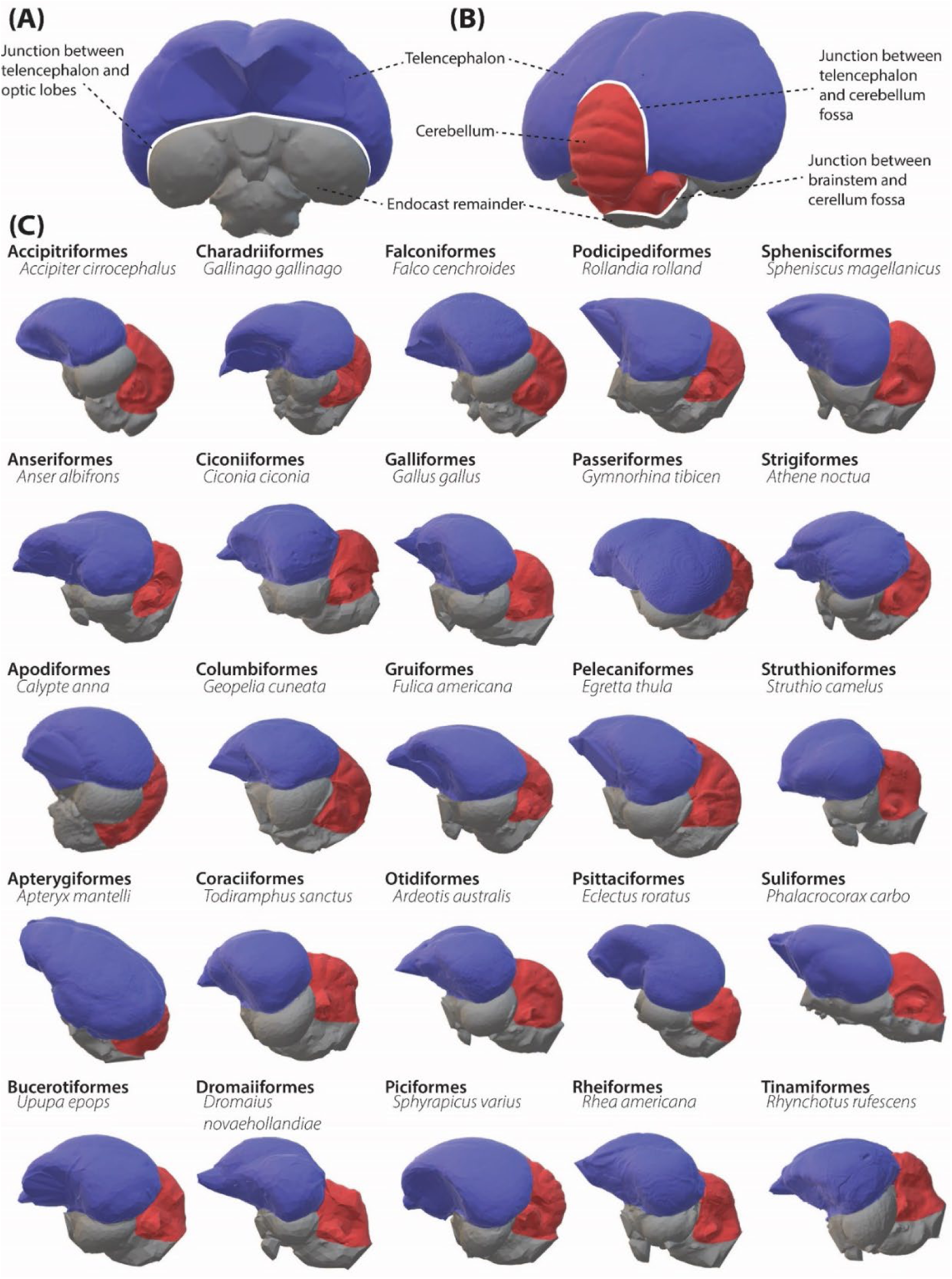
The endocast of a Eurasian treecreeper (*Certhia familiaris*) in rostral (A) and caudal views (B). Both images show osteological landmarks used to delineate the endocast telencephalon and cerebellum for measurements. The telencephalon is blue, the cerebellum is red, and the rest of the endocast is grey. (C) Endocasts of a representative species of each of the orders examined in this study using the same colour coding as in A and B. Note that endocasts shown here are of reduced resolution and are not to scale.

### Statistical analysis

All data were natural log transformed for analyses. To assess if the absolute size of the endocast surface areas is an appropriate proxy of the absolute size of their brain counterpart, we completed ordinary least squares (OLS) regression analyses with brain region volume as predictor and endocast surface area as response variables in R [60]. From these, we calculated confidence intervals (the confidence of the fit), prediction intervals (the confidence with which an unknown specimen could be given a value), and the line of best fit for each set of measurements: whole endocast, telencephalon and cerebellum. We expected the endocast surface areas to scale isometrically with the brain volume, with surface area scaling at two-thirds the volume (i.e., yielding slopes of 0.67). To test this, we assessed the slope coefficients of the linear relationships and additionally used Wald tests in the *car* package [62], which uses the model coefficients and coefficient-covariance matrix to determine if the relationship significantly deviated from the isometric slope of 0.67 [63]. Note that the relationship between brains and endocasts is not expected to be impacted by phylogenetic structure of the data. For this reason, linear modelling was the preferred statistical procedure. However, phylogenetic generalised least squares models were also computed using the *evomap* package [64] to assess if phylogenetic relationships impact on the estimates of the slope [65].

As we have previously shown that the variation in voxel sizes of the CT scans may cause minor discrepancies for endocast measurements [8] we tested if this may have had an artificial effect on endocast size. We did this by calculating the residuals of OLS regression analyses using the whole endocast surface area and whole brain volume as dependent variables and voxel size as the predictor variable. We expected that if this factor does not influence the results that the residuals should show a similar relationship to our comparisons of the endocast and brain measurements.

We also computed the relative size for both the telencephalon and cerebellum by scaling them with the remainder of the brain as the dependent variable [66], both in the surface area (endocast-based) data and the volume (histology-based) data. This was to confirm that the slopes of the scaling relationships were comparable between the two data sets. Because phylogenetic relationships exert a significant effect on the relative size of brain regions [43, 65, 67, 68], we used both linear modelling and phylogenetic generalised least squares modelling to derive the slopes and intercepts.

We derived our phylogeny by pruning the family-level phylogeny by Stiller, Feng [69] and used Mesquite [70] to place species on this tree using order specific phylogenetic studies [71–81]. We then completed the PGLS regression analyses and calculated phylogenetic confidence intervals (CI) for each of the regions against the remainder of either the endocast or brain using the *evomap* package [64].

## RESULTS

Our comparisons of the surface area measurements for the whole endocast, telencephalon, and cerebellum against their brain volume counterparts yielded highly significant associations (Figure 2; Table 1) which are unlikely to be influenced by the voxel size (see electronic supplementary material, figure S2). Furthermore, the resulting slopes indicate isometric relationships (Table 1) which are not significantly different from the expected 0.67 according to the Wald test comparison (Table 1).While this overall pattern is highly consistent, some species fell outside of the prediction intervals (PI) for the whole sample, meaning that their surface area measurements were not expected given their corresponding volumes. In total, nine species fell outside of the PIs for at least one of the regressions with three outside of the PIs for all regressions: the satin bowerbird (*Ptilonorhynchus violaceus*; Passeriformes), black-capped chickadee (*Poecile atricapillus*; Passeriformes), and the collared sparrowhawk (*Accipiter cirrocephalus*; Accipitriformes) (Figure 2).

**Figure 2.**
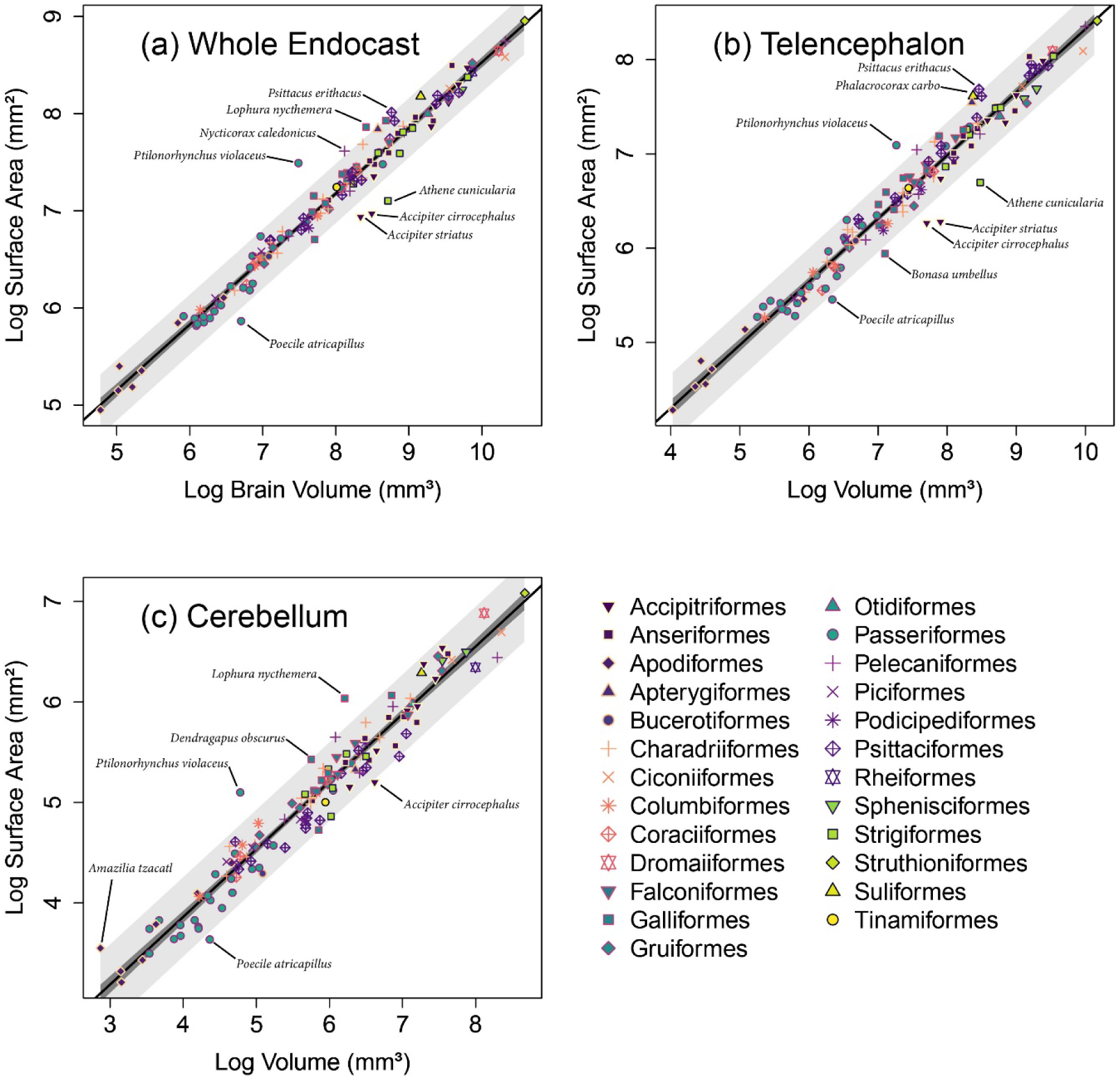
Log transformed measurements of the (a) whole endocast, (b) telencephalon, and (c) cerebellum surface areas measured from endocasts regressed against their brain volume counterparts for 136 species of birds. The orders from this data are represented by unique combinations of colours with point shapes. The black line represents the ordinary least-squares linear model regression; dark grey area represents 95% confidence intervals; light grey area represents 95% prediction intervals.

**Table 1.** Results from ordinary least squares (OLS) regression analyses and Wald tests. OLS results show the significant (p=<0.05) relationships between the endocast surface area measurements and the brain volume measurements. Wald test results show that slopes do not significantly (*p*=>0.05) differ from the expected isometric relationship between surface area and volume measurements (0.67).

| Dependent | Predictor | *df* | F-statistic | *p*-value | *y* intercept | R<sup>2</sup> | Slope | Wald test *p*-value |
| --- | --- | --- | --- | --- | --- | --- | --- | --- |
| log Endo. SA | log Brain VOL | 134 | 3969 | <0.01 | 1.78 | 0.97 | 0.68 | 0.64 |
| log Tele. SA | log Tele. VOL | 132 | 2954 | <0.01 | 1.62 | 0.96 | 0.67 | 0.99 |
| log Cere. SA | log Cere. VOL | 132 | 2324 | <0.01 | 1.17 | 0.95 | 0.67 | 0.84 |

Abbreviations: SA, surface area; Vol., volume; Endo., endocast; Tele., telencephalon; Cere., cerebellum.

For the comparisons of the relative sizes of the telencephalon and cerebellum, the surface area for these regions regressed against the endocast remainder yielded similar slopes to that of the region volume regressed against the brain remainder, for both the OLS and PGLS (Figure 3; Table 2). Parrots (Psittaciformes) appear to deviate more from the regression lines in Figure 3 for both regions. This is because parrots have relatively large telencephala and small cerebella compared with other species [43]. The surface area measurements show a somewhat larger deviation for the telencephalon, which could arise from the relationship between surface area and volume or that the optic lobes are smaller in parrots [43, 82], allowing the telencephalon to spread out more. One species, the Kākāpō (*Strigops habroptilus*), is outside of the PGLS confidence interval for the expected range of the telencephalon surface area scaled against the endocast remainder surface area (Figure 3a). This species is a uniquely nocturnal and flightless parrot that is neuroanatomically distinct within its clade [54] which may then result in an atypical relationship between endocast and brain measurements. Overall, these results indicate that the scaling relationship for both brain regions is similar when measured from endocast surface areas and brain volumes.

**Figure 3.**
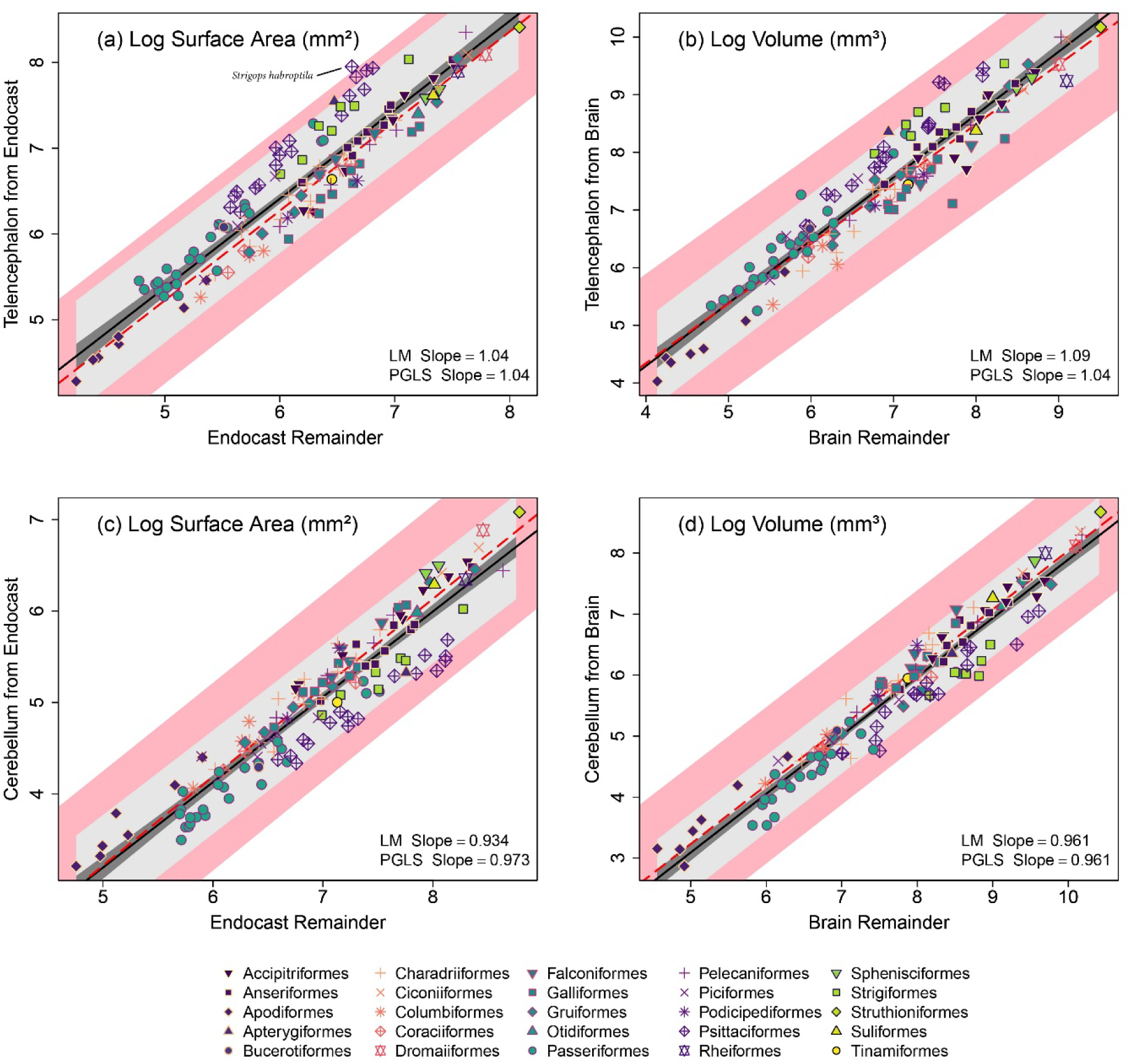
Log transformed measurements comparing the scaling relationships of the telencephalon for (a) the surface area against endocast remainder, and (b) the volume against brain remainder, as well as the cerebellum for (c) the surface area against endocast remainder, and (d) the volume against brain remainder for 136 species of birds. The orders from this data are represented by unique combinations of colours with point shapes. Ordinary least squares regression (LM slope) is shown by the black line; dark grey for the 95% confidence intervals; light grey for the 95% prediction intervals. Phylogenetic generalised least squares (PGLS) regression shown by the dashed red line; pink area shows the 95% confidence intervals of the PGLS.

**Table 2.** Results from phylogenetic generalised least squares (PGLS) regression models used to compare the scaling relationship of endocast surface area and brain volume measurements of the telencephalon and cerebellum.

| Dependent | Predictor | *df* | *t*-value | *p*-value | *y* intercept | Slope |
| --- | --- | --- | --- | --- | --- | --- |
| Tele. SA. | Endo. Remainder | 134 | 32.2 | <0.01 | 0 | 1.04 |
| Tele. VOL | Brain remainder | 134 | 28.97 | <0.01 | 0.2 | 1.04 |
| Cere. SA | Endo. Remainder | 134 | 29.35 | <0.01 | -1.65 | 0.97 |
| Cere. VOL | Brain remainder | 134 | 33.48 | <0.01 | -1.57 | 0.96 |

Abbreviations: SA, surface area; Vol., volume; Endo., endocast; Tele., telencephalon; Cere., cerebellum.

## DISCUSSION

Our results reveal that avian endocast surface areas for the whole endocast, the telencephalon, and the cerebellum exhibit tight isometric relationships with the volume of their brain counterparts. This result is in spite of a sample size of one individual per species for the endocast measurements and a reliance on different individuals for the endocast and brain measurements, both of which could have reduced the strength of the association between the two datasets [31]. By including a broad range of species, our validation supports the use of endocast measurements for these regions in future studies, while also supporting the conclusions from previous studies which relied on such measurements [8, 9, 25, 36, 37, 83, 84]. This result is particularly beneficial for studies on rare species [8, 25], and palaeontological specimens [9, 85] for which brain measurements are not attainable.

We also found that the relative sizes for both the telencephalon and cerebellum regions are consistent between the endocast surface areas and the soft tissue brain volumes, supporting their use in comparisons of brain composition [43]. This is in spite of the absence of the entire anterior lobe of the cerebellum in endocasts which is generally hidden from external view by the telencephalon, and the cerebellar surface often obscured by the occipital sinus [3, 32, 86, 87]. For other endocast validation research, such as that of the optic lobes and wulst [28], the regions are typically clearly defined and largely unobscured. Nonetheless, our findings strongly suggest that the portion of the cerebellum visible in endocasts is sufficient for estimating its relative volume.

While there is a strong brain–endocast association despite our reliance on different individuals, some species showed disparities between specimens, likely due to intraspecific variation in brain size and composition arising from sexual dimorphism [88–90] or subspecific, clinal, and other regional variation [91]. In fact, in a previous study, we found substantial skull and endocast size differences for two collared sparrowhawk specimens, likely due to sexual dimorphism [25]. Within this study, we suspect variation from sexual dimorphism in hawks [88, 90], and pheasants [56, 92],and regional variation for the bowerbird [89], chickadee [93], burrowing owl [94] and cormorant [95]. Furthermore, as variation occurred when comparing volumes and surface areas directly to one another but did not appear to be present in the analyses of relative brain region size we suspect this variation stems from using different specimens for both datasets. While variation is expected, this raises concerns regarding the sample size needed to represent a species in comparative studies, particularly of narrow taxonomic scope. We nevertheless note the high correspondence between the brain and endocast datasets, suggesting that these measurements can generally be used with small sample sizes.

The telencephalon and cerebellum make up the majority of the brain and endocast surface, making their inclusion necessary for studies on brain composition [43]. By using known brain–behaviour associations, such as the telencephalon size with cognitive abilities [36, 40, 96], and the cerebellum with flight abilities [37, 38, 42], some aspects of ecology and behaviours may be inferred from these regions [40, 46, 97–99]. Furthermore, even though some behaviourally relevant brain regions (such as the hippocampus [100]) will always remain hidden within the brain, holistic approaches to endocast comparisons can improve the accuracy of inferences by accounting for the influences that changes in brain regions sizes can have on one another [14, 43, 99, 101]. For example, the relative size of the telencephalon in owls is comparable with that of parrots [96], but this is largely due to their hypertrophied wulst in relation to vision [43] and not the regions of the telencephalon associated with cognition [102].

Overall, this study has confirmed the strong relationships between the surface areas of endocasts and brain volumes in birds for absolute and relative size, both wholly and when divided into the telencephalon and cerebellum regions. This result is in concordance with other endocast validations [28, 31], and we suggest that further expanding the use of avian endocasts is plausible. However, these methods remain speculative beyond the crown group Aves, such as for non-avian dinosaurs [2, 4, 29]. Outside of validating bird endocasts, comparisons with other living archosaurs, the crocodilians, may assist with resolving the brain–endocast correspondence [2] and provide further evidence for the strengths and limitations of these methods [4, 30]. With the increased accessibility that CT scanning and associated digital reconstruction methods present [1, 10, 103], expanding and refining endocast uses has the potential to shed light on the lives of extinct species and further resolve our understanding of vertebrate brain evolution.

## Supporting information

electronic supplementary material

## Acknowledgements

We thank Maya Penck (South Australian Museum) and Karen Roberts (Museum Victoria) for access to specimens scanned for this study. Thanks to Jay Black at the University of Melbourne, along with Sophie Rapagna and Egon Perilli at Flinders Microscopy, for completing the scans of these specimens. We thank Prof. Roger Benson and the team behind the openVertebrate (oVert) project for provision of CT scans which made this validation possible. We thank the MorphoSource team as well as those who contribute to this repository. This study was mostly conducted on the traditional lands of the Kaurna (Adelaide, South Australia) and Niitsitapi (Lethbridge, Alberta) people.

