## Supplementary material for "Avian telencephalon and cerebellum volumes can be accurately estimated from digital brain endocasts": electronic supplementary material

**Detailed explanation of endocast borders**

The posterior border of the telencephalon fossa was defined as the junction where it meets the cerebellar fossa and continues ventrally through the crista tentorialis above the fossa tecti mesenchephali. These borders then meet rostrally above the optic chiasma. For the cerebellum fossa, the dorsal border was defined by the junction that meets the telencephalon fossa and continues along the caudal border of the fossa tecti mesencephali on either side. This border continues as the crista present at the ventral region of the cerebellum fossa, dorsal to the brainstem fossa, meeting in the posterior region of the foramen magnum.

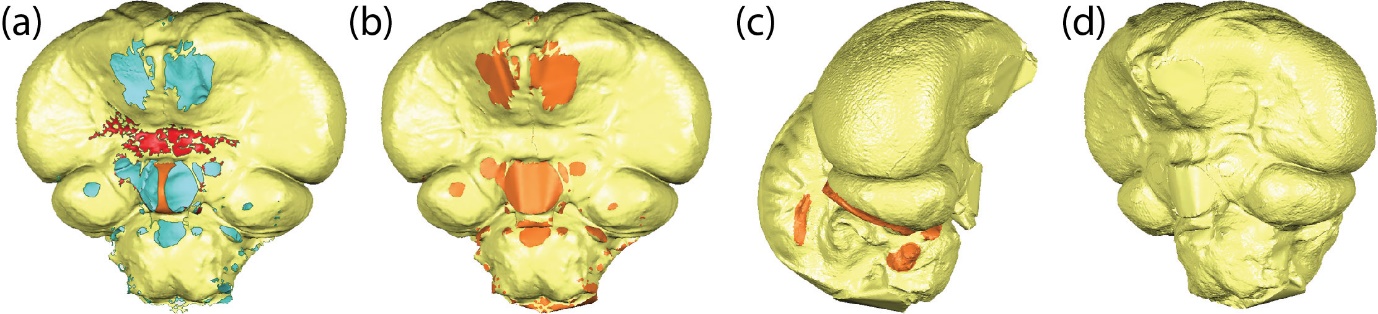
 **Supplementary Figure 1.** Endocast of Collared Sparrowhawk (*Accipiter cirrocephalus*) when (a) imported into Geomagic from endomaker mesh (yellow: outer faces, red and orange: selected excess material for removal, blue: inner faces), (b) holes filled with “flat” fill setting, (c) nerves and excess material (ie. blood vessels) selected for removal, and (d) complete endocast.

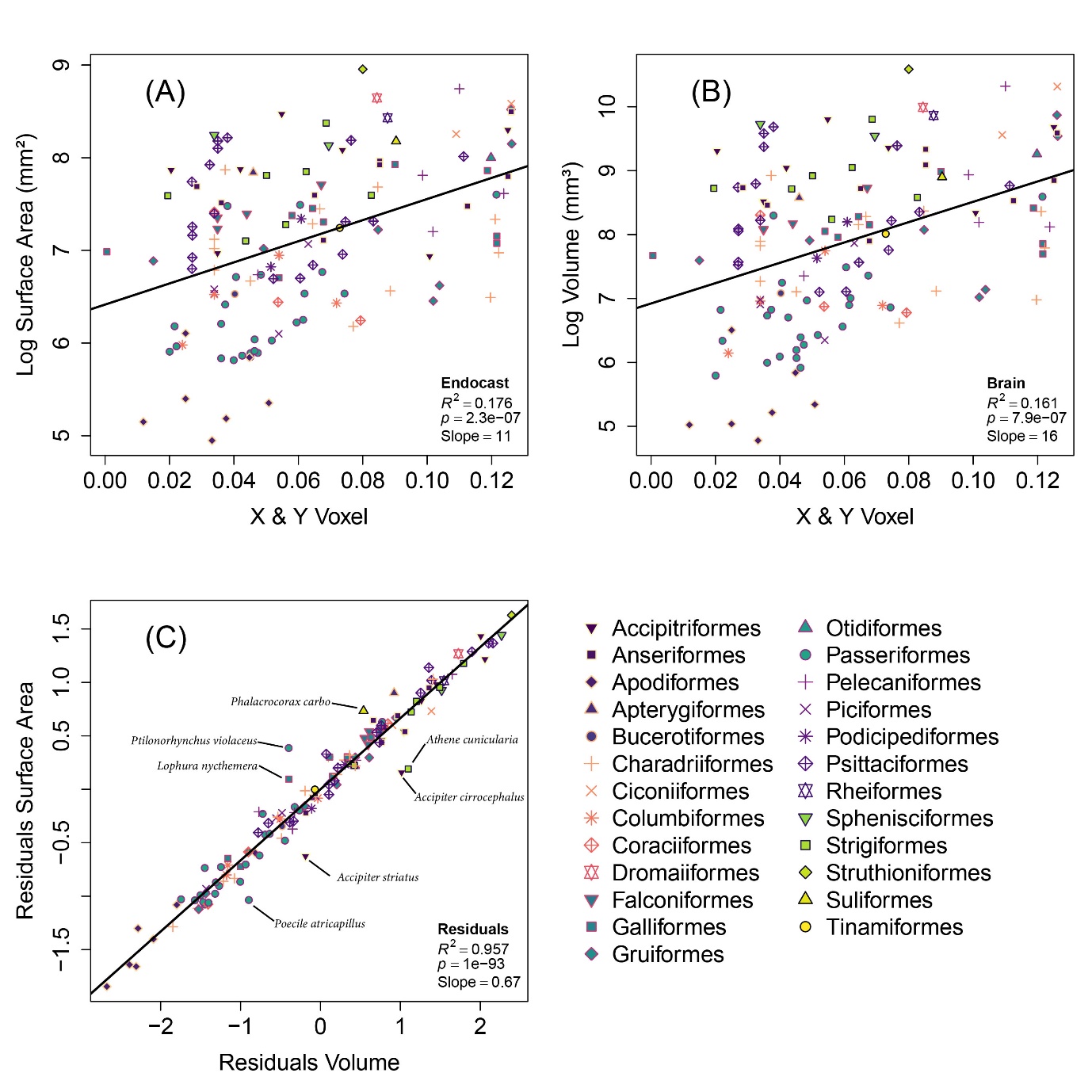

**Supplementary Figure 2.** Log transformed measurements of the (a) endocast surface area and (b) brain volume regressed against the X & Y voxel dimensions for each species. Ordinary least-squares regression analysis was used to calculate the residuals for each species which are plotted against one another in (c) as well as the r-squared, p-value and slope shown in the bottom right corner for each plot.
